# Single-Case Disconnectome Lesion-symptom Mapping: Identifying Two Subtypes of Limb Apraxia

**DOI:** 10.1101/2022.02.22.481524

**Authors:** Rachel Metzgar, Harrison Stoll, Scott T. Grafton, Laurel J. Buxbaum, Frank E. Garcea

**Author notes:** **Corresponding Author:** Frank E. Garcea, Department of Neurosurgery, University of Rochester Medical Center, Rochester, NY 14642.

## Abstract

Influential theories of skilled action posit that distinct cognitive mechanisms and neuroanatomic substrates support meaningless gesture imitation and tool use pantomiming, and poor performance on these tasks are hallmarks of limb apraxia. Yet prior research has primarily investigated brain-behavior relations at the group level; thus, it is unclear whether we can identify individuals with isolated impairments in meaningless gesture imitation or tool use pantomiming whose performance is associated with a distinct neuroanatomic lesion profile. The goal of this study was to test the hypothesis that individuals with disproportionately worse performance in meaningless gesture imitation would exhibit cortical damage and white matter disconnection in left fronto-parietal brain regions, whereas individuals with disproportionately worse performance in tool use pantomiming would exhibit cortical damage and white matter disconnection in left temporo-parietal brain regions. Fifty-eight participants who experienced a left cerebrovascular accident took part in a meaningless gesture imitation task, a tool use pantomiming task, and a T1 structural MRI. Two participants were identified who had relatively small lesions and disproportionate impairments on one task relative to the other, as well as below-control-level performance on one task and not the other. Using these criteria, one participant was disproportionately impaired at meaningless gesture imitation, and the other participant was disproportionately impaired at pantomiming tool use. Graph theoretic analysis of each participant’s structural disconnectome demonstrated that disproportionately worse meaningless gesture imitation performance was associated with disconnection among the left inferior parietal lobule, the left superior parietal lobule, and the left middle and superior frontal gyri, whereas disproportionately worse tool use pantomiming performance was associated with disconnection between left temporal and parietal regions. Our results demonstrate that relatively focal lesions to specific portions of the Tool Use Network can be associated with distinct limb apraxia subtypes.

**Highlights:**

- Distinct subtypes of apraxia are supported by distinct neuroanatomic substrates.
- Fronto-parietal lesions are associated with meaningless imitation deficits.
- Temporo-parietal lesions are associated with tool use pantomiming deficits.
- Structural connectivity among cortical areas supports a distributed praxis system.

## 1.1 Introduction

Limb apraxia is a disorder of skilled action production that is associated with lesions to the left inferior parietal lobule. Despite relatively intact sensory and motor function, patients with limb apraxia (hereafter, apraxia) can be impaired when asked to imitate meaningless gestures or pantomime tool use gestures from command (Buxbaum & Saffran, 2002; Garcea et al., 2013; Goldenberg, 2009; Johnson-Frey, 2004; Leiguarda & Marsden, 2000; Rothi et al., 1991). Historically, cognitive models of apraxia posit a distinction between production and conceptual mechanisms supporting praxis. Patients with ideomotor apraxia have a deficit in the transformation of action knowledge into a sensory-motor format for action production. In contrast, patients with ideational apraxia have a deficit in retrieving conceptual knowledge of actions, which affects the selection and production of skilled actions when performing compound actions (e.g., see Buxbaum, 2001; Cubelli et al., 2000; De Renzi & Lucchelli, 1988; Liepmann, 1905; Ochipa et al., 1989; Roy & Square, 1985; Schwartz et al., 2002; Wheaton & Hallett, 2007). Ideational and ideomotor subtypes of apraxia have not been linked to distinct neuroanatomic substrates.

Neuropsychological dissociations between meaningless gesture imitation and tool use pantomime suggest that distinct cognitive mechanisms support the imitation and pantomiming of gestures (e.g., see Buxbaum et al., 2014; Tessari et al., 2007). In this project, we focused our analyses on individual cases with relatively small left cerebrovascular accident (LCVA) lesions, as this approach permitted a fine-grained neuroanatomic investigation of the lesion sites and stroke-induced structural disconnection in individuals who were disproportionately impaired when pantomiming tool use (PTU) relative to meaningless gesture imitation (MGI), and vice versa.

Investigations of the neuroanatomy supporting PTU and MGI have focused on visuomotor computations associated with the dorsal visual pathway (Buxbaum et al., 2003; Chao & Martin, 2000; Goldenberg, 2014; Goodale et al., 1991; Orban & Caruana, 2014). Motivated in part from research in non-human primates (e.g., see Rizzolatti & Matelli, 2003), the 2 Action Systems Plus (2AS+) neurocognitive model distinguishes between a dorso-dorsal and a ventro-dorsal pathway for action within the human dorsal visual pathway (see Binkofski & Buxbaum, 2013). The dorso-dorsal pathway supports the buffering, sequencing, and transformation of current visual information into a sensory-motor format for reaching and grasping, supported by the bilateral superior parietal lobule, bilateral intraparietal sulci, and dorsal premotor cortex. The ventro-dorsal pathway transforms action knowledge (stored in part in the left posterior middle temporal lobe) into a sensory-motor format for skilled tool manipulation in the service of a behavioral goal, supported by the left supramarginal gyrus, left ventral premotor cortex, and left inferior frontal gyrus (for discussion, see Buxbaum, 2017; Gallivan & Culham, 2015; Hoeren et al., 2014; Lingnau & Downing, 2015; Mahon & Caramazza, 2011; Martin, 2007; Vry et al., 2015).

At the group level, the lesion loci associated with impairments in MGI and PTU are consistent with the predictions of the 2AS+ model. For instance, voxel-based lesion-symptom mapping (VLSM) investigations demonstrate that errors in PTU are associated with lesions to temporo-parietal structures, including the left supramarginal gyrus and left posterior middle temporal gyrus (Buxbaum et al., 2014; Martin et al., 2016; Sperber et al., 2019; Weiss et al., 2016), whereas errors in MGI are associated with lesions to the left inferior parietal lobule, left superior parietal lobule, left pre- and post-central gyri, and left dorsal premotor cortex (Buxbaum et al., 2014; Haaland et al., 2000; Hoeren et al., 2014; Mengotti et al., 2013; Tessari et al., 2007; Weiss et al., 2001; Wong et al., 2019). Although the lesion loci of MGI and PTU partially overlap in frontal, parietal, and temporal nodes of the left hemisphere, a consistent finding across studies is that MGI errors are associated with fronto-parietal lesion sites (for discussion, see Rumiati et al., 2009), whereas PTU errors are associated with temporo-parietal lesion sites (for recent discussion, see Buxbaum & Randerath, 2018).

Recent neuroimaging studies have demonstrated that, in addition to lesion location, structural and functional connectivity disruption among temporal, frontal, and parietal nodes that comprise a ‘Tool Use Network’ may be associated with apraxia severity. For example, Watson and colleagues (2019) found that in a group of LCVA participants, lower tool use pantomiming accuracy was associated with reduced resting state functional connectivity between the left inferior parietal lobule and left middle temporal gyrus (see also Garcea, Almeida, et al., 2019). Garcea and colleagues (2020) showed that after controlling for shared variance with MGI performance, reduced PTU performance was associated with the degree of white matter structural disconnection between the left inferior parietal lobule and left middle temporal gyrus, and between the left inferior parietal lobule and left middle frontal gyrus (see also Bi et al., 2015). These findings suggest that distinct aspects of praxis are supported by distinct cortical nodes and white matter pathways. However, prior group-level VLSM and connectome-based lesion-symptom mapping studies cannot determine whether individual LCVA participants show distinct subtypes of apraxia associated with distinct neuroanatomic substrates.

The focus of the current project was to investigate the lesion sites and structural disconnectivity of individual LCVA participants who had relatively small lesions and who exhibited disproportionately worse performance between PTU and MGI tasks. Our goal was to identify participants with distinct conceptual or production apraxia subtypes, and to test whether those subtypes were associated with distinct neuroanatomical lesion sites and structural disconnectivity patterns. Testing for differences in individual case performance permits a test of two hypotheses: (1) Individuals with disproportionately worse performance in MGI relative to PTU will exhibit cortical damage and white matter disconnection in left fronto-parietal regions, and (2) Individuals with disproportionately worse performance in PTU relative to MGI will exhibit cortical damage and white matter disconnection in left temporo-parietal regions.

### 2.1 Materials and Methods

#### 2.1.1 LCVA Participants

This study is a re-analysis of group-level data previously reported by Garcea and colleagues (2020). Sixty-six chronic LCVA participants were recruited from the Neuro-Cognitive Research Registry at Moss Rehabilitation Research Institute (Schwartz et al., 2005). Of those participants, 58 completed the tool use pantomiming task and the meaningless imitation task and had undergone a research-quality T1 structural MRI. We restricted our analysis to 29 LCVA participants with relatively smaller lesions using a median split of total lesion volume (13 female; mean age = 58.10 years, SD = 11.13 years, range = 31 – 80 years; mean education = 14.17 years, SD = 3.07 years, range = 9 – 21 years). See Supplemental Table 1 for demographic and behavioral data of the 29 LCVA participants with larger lesions. Participants were right-hand dominant (1 reported as ambidextrous) and had suffered a single left hemisphere stroke at least 3 months prior to testing (mean number of months since stroke = 29.41 months, SD = 29.42 months; range = 4 – 114 months). Participants were excluded if they had a history of psychosis, drug or alcohol abuse, co-morbid neurological disorder, or severe language comprehension deficits indicated by a score below 4 on the comprehension subtest of the Western Aphasia Battery (Kertesz, 1982). See Table 1 for demographic information and Figure 1 for the maximally disconnected structural disconnectome for each LCVA participant. Behavioral data analysis code can be downloaded here: https://github.com/frankgarcea/MetzgarColleagues.git

**Figure 1.**
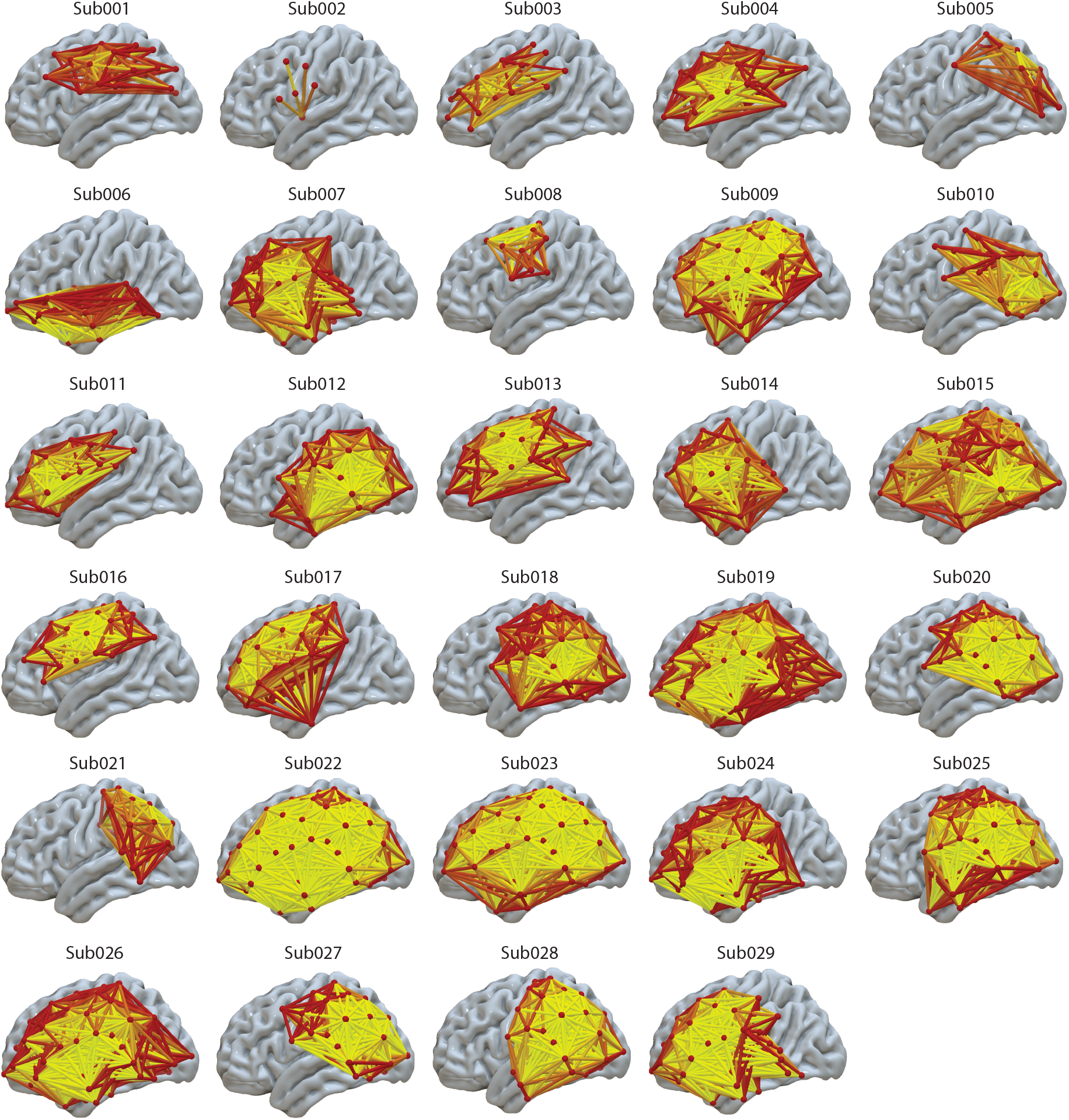
Left hemisphere rendering of each LCVA participant’s maximally disconnected subgraph. Each participant’s maximally disconnected subgraph is presented on the ICBM152 template. Each node (red sphere) is positioned at the center of mass of each Lausanne2008 atlas region of interest. Yellow edges exhibit greater disconnectivity relative to red edges.

**Table 1.**
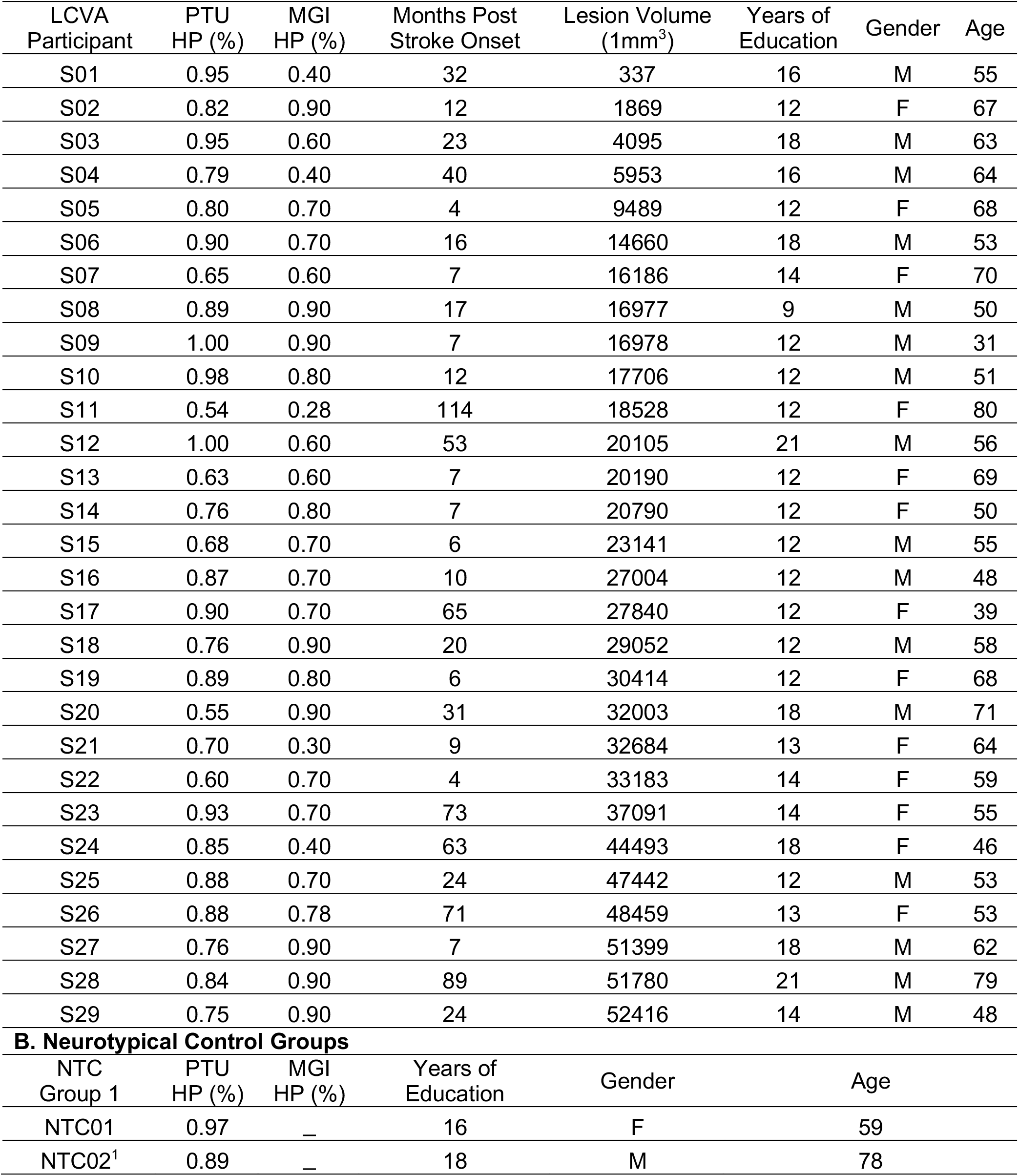
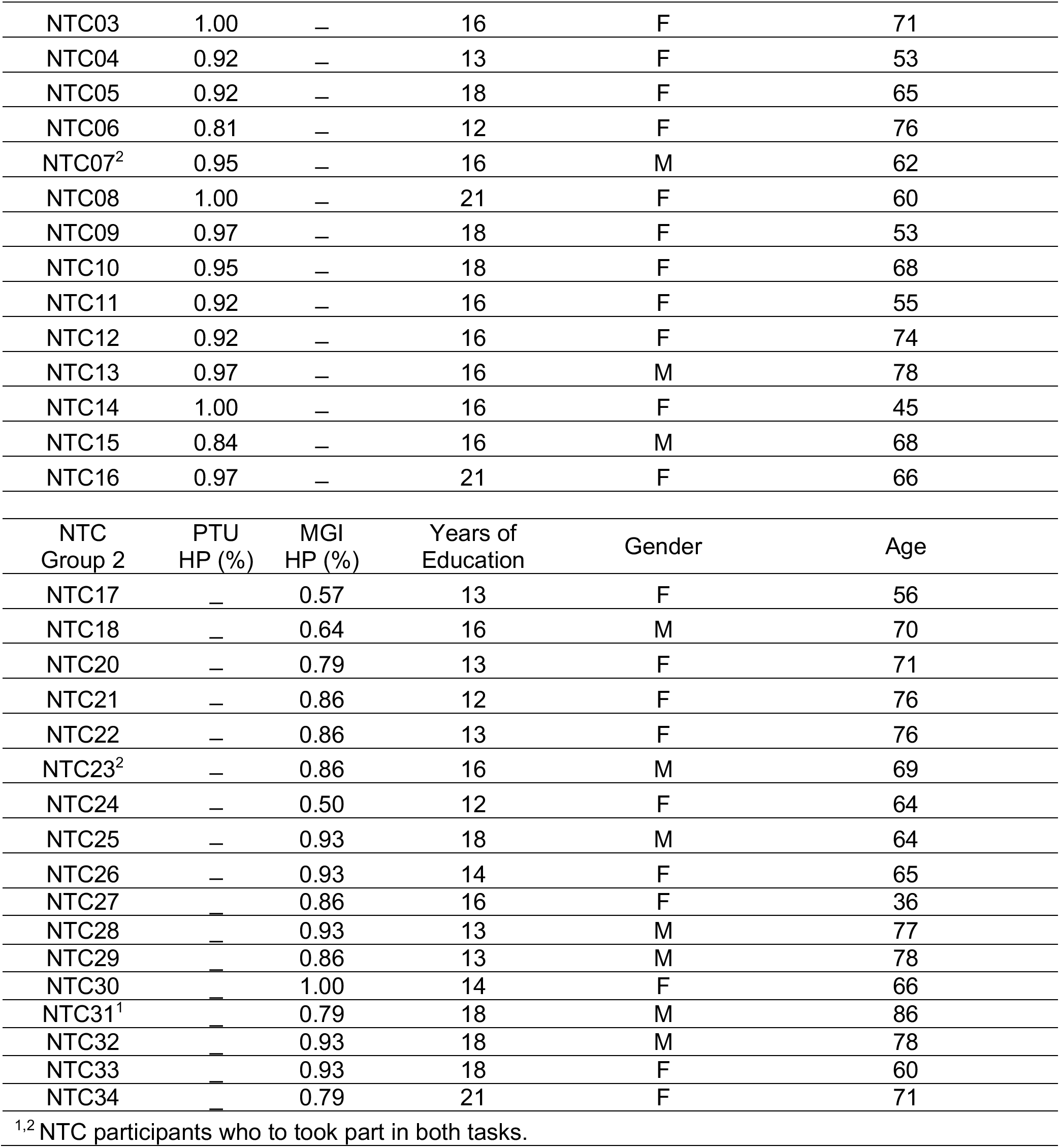
Demographic information and lesion volume for each LCVA participant. *Abbreviations*. PTU, Pantomime of tool use to the sight of objects hand posture accuracy; MGI, meaningless gesture imitation hand posture accuracy.

| LCVA Participant | PTU HP (%) | MGI HP (%) | Months Post Stroke Onset | Lesion Volume (1mm<sup>3</sup>) | Years of Education | Gender | Age |
| --- | --- | --- | --- | --- | --- | --- | --- |
| S01 | 0.95 | 0.40 | 32 | 337 | 16 | M | 55 |
| S02 | 0.82 | 0.90 | 12 | 1869 | 12 | F | 67 |
| S03 | 0.95 | 0.60 | 23 | 4095 | 18 | M | 63 |
| S04 | 0.79 | 0.40 | 40 | 5953 | 16 | M | 64 |
| S05 | 0.80 | 0.70 | 4 | 9489 | 12 | F | 68 |
| S06 | 0.90 | 0.70 | 16 | 14660 | 18 | M | 53 |
| S07 | 0.65 | 0.60 | 7 | 16186 | 14 | F | 70 |
| S08 | 0.89 | 0.90 | 17 | 16977 | 9 | M | 50 |
| S09 | 1.00 | 0.90 | 7 | 16978 | 12 | M | 31 |
| S10 | 0.98 | 0.80 | 12 | 17706 | 12 | M | 51 |
| S11 | 0.54 | 0.28 | 114 | 18528 | 12 | F | 80 |
| S12 | 1.00 | 0.60 | 53 | 20105 | 21 | M | 56 |
| S13 | 0.63 | 0.60 | 7 | 20190 | 12 | F | 69 |
| S14 | 0.76 | 0.80 | 7 | 20790 | 12 | F | 50 |
| S15 | 0.68 | 0.70 | 6 | 23141 | 12 | M | 55 |
| S16 | 0.87 | 0.70 | 10 | 27004 | 12 | M | 48 |
| S17 | 0.90 | 0.70 | 65 | 27840 | 12 | F | 39 |
| S18 | 0.76 | 0.90 | 20 | 29052 | 12 | M | 58 |
| S19 | 0.89 | 0.80 | 6 | 30414 | 12 | F | 68 |
| S20 | 0.55 | 0.90 | 31 | 32003 | 18 | M | 71 |
| S21 | 0.70 | 0.30 | 9 | 32684 | 13 | F | 64 |
| S22 | 0.60 | 0.70 | 4 | 33183 | 14 | F | 59 |
| S23 | 0.93 | 0.70 | 73 | 37091 | 14 | F | 55 |
| S24 | 0.85 | 0.40 | 63 | 44493 | 18 | F | 46 |
| S25 | 0.88 | 0.70 | 24 | 47442 | 12 | M | 53 |
| S26 | 0.88 | 0.78 | 71 | 48459 | 13 | F | 53 |
| S27 | 0.76 | 0.90 | 7 | 51399 | 18 | M | 62 |
| S28 | 0.84 | 0.90 | 89 | 51780 | 21 | M | 79 |
| S29 | 0.75 | 0.90 | 24 | 52416 | 14 | M | 48 |
| **B. Neurotypical Control Groups** |  |  |  |  |  |  |  |
| NTC Group 1 | PTU HP (%) | MGI HP (%) | Years of Education | Gender | Age |  |  |
| NTC01 | 0.97 | — | 16 | F | 59 |  |  |
| NTC02<sup>1</sup> | 0.89 | — | 18 | M | 78 |  |  |

| NTC03 | 1.00 | — | 16 | F | 71 |
| --- | --- | --- | --- | --- | --- |
| NTC04 | 0.92 | — | 13 | F | 53 |
| NTC05 | 0.92 | — | 18 | F | 65 |
| NTC06 | 0.81 | — | 12 | F | 76 |
| NTC07<sup>2</sup> | 0.95 | — | 16 | M | 62 |
| NTC08 | 1.00 | — | 21 | F | 60 |
| NTC09 | 0.97 | — | 18 | F | 53 |
| NTC10 | 0.95 | — | 18 | F | 68 |
| NTC11 | 0.92 | — | 16 | F | 55 |
| NTC12 | 0.92 | — | 16 | F | 74 |
| NTC13 | 0.97 | — | 16 | M | 78 |
| NTC14 | 1.00 | — | 16 | F | 45 |
| NTC15 | 0.84 | — | 16 | M | 68 |
| NTC16 | 0.97 | — | 21 | F | 66 |
| NTC<br>Group 2 | PTU<br>HP (%) | MGI<br>HP (%) | Years of<br>Education | Gender | Age |
| NTC17 | — | 0.57 | 13 | F | 56 |
| NTC18 | — | 0.64 | 16 | M | 70 |
| NTC20 | — | 0.79 | 13 | F | 71 |
| NTC21 | — | 0.86 | 12 | F | 76 |
| NTC22 | — | 0.86 | 13 | F | 76 |
| NTC23<sup>2</sup> | — | 0.86 | 16 | M | 69 |
| NTC24 | — | 0.50 | 12 | F | 64 |
| NTC25 | — | 0.93 | 18 | M | 64 |
| NTC26 | — | 0.93 | 14 | F | 65 |
| NTC27 | — | 0.86 | 16 | F | 36 |
| NTC28 | — | 0.93 | 13 | M | 77 |
| NTC29 | — | 0.86 | 13 | M | 78 |
| NTC30 | — | 1.00 | 14 | F | 66 |
| NTC31<sup>1</sup> | — | 0.79 | 18 | M | 86 |
| NTC32 | — | 0.93 | 18 | M | 78 |
| NTC33 | — | 0.93 | 18 | F | 60 |
| NTC34 | — | 0.79 | 21 | F | 71 |

<sup>1,2</sup> NTC participants who took part in both tasks.

#### 2.1.2 Neurotypical Control Participants

Because of COVID-19 testing restrictions, we were unable to acquire PTU and MGI performance in the same cohort of neurotypical control (NTC) participants; thus, we are reporting data from two previous studies. Together, there were thirty-one participants recruited from the Neuro-Cognitive Research Registry at Moss Rehabilitation Research Institute. Participants were excluded if they had a history of psychosis, drug or alcohol abuse, or neurological disorder (see Table 1). The NTC groups were not different in terms of age (t < 1) and education (t(29) = 1.19, p = 0.24).

#### 2.1.3 Neurotypical Control (NTC) Group 1

As reported previously (Watson and Buxbaum (2015), 16 right-hand dominant NTC participants (12 female) performed a tool use pantomiming task (mean age = 64.43 years, SD = 9.80 years; mean education = 16.90 years, SD = 2.40 years).

#### 2.1.4 Neurotypical Control (NTC) Group 2

Seventeen NTC participants (7 female) performed a meaningless gesture imitation task (unpublished data). Fifteen NTC Group 2 participants were right-hand dominant and 2 were ambidextrous (mean age = 68.42 years, SD = 11.15 years; mean education = 15.18 years, SD = 2.70 years).

### 2.2 Neuroimaging Acquisition

#### 2.2.1 Acquisition of Anatomic Scans

MRI scans included whole-brain T1-weighted structural MRI collected on a 3T (Siemens Trio, Erlangen, Germany; repetition time = 1620 ms, echo time = 3.87 milliseconds, field of view = 192 x 256 mm, 1×1×1 mm voxels) or a 1.5T (Siemens Sonata, repetition time = 3,000 milliseconds, echo time = 3.54 milliseconds, field of view = 24 cm, 1.25×1.25×1.25 mm voxels) scanner using an eight-channel or sixty-four channel head coil. Lesions were manually segmented on each LCVA participant’s high-resolution T1-weighted structural images. Lesioned voxels, consisting of both grey and white matter, were assigned a value of 1 and preserved voxels were assigned a value of 0. Binarized lesion masks were then registered to a standard template (Montreal Neurological Institute “Colin27”) using a symmetric diffeomorphic registration algorithm (Avants et al., 2008; www.picsl.upenn.edu/ANTS). Volumes were first registered to an intermediate template comprised of healthy brain images acquired on the same scanner. Volumes were then mapped onto the “Colin27” template to complete the transformation into standardized space. To ensure accuracy during the transformation process, lesion maps were subsequently inspected by a neurologist (H.B. Coslett) naive to the behavioral data of the study. For increased accuracy, the pitch of the template was rotated to approximate the slice plane of each LCVA participant’s scan. This method has been demonstrated to achieve high intra- and inter-rater reliability (e.g., see Schnur et al., 2009). See Supplemental Figure 1 for a rendering of each LCVA participant’s lesion.

### 2.3 Neuropsychological Testing of Tool Use Pantomiming and Meaningless Imitation

#### 2.3.1 Pantomiming Tool Use (PTU)

The PTU task included 40 photographs of manipulable objects (tools) taken from the BOSS database (Brodeur et al., 2010). Tools included items with distinct use actions, including construction tools (e.g., wrench), household articles (e.g., teapot), office supplies (e.g., scissors), and bathroom items (e.g., razor). Fourteen of the tool use actions were directed towards the body (e.g., cup, fork), and 26 actions were directed away from the body (e.g., hammer, scissors). Each trial of the task began with the presentation of a 400-by-400 pixel color photograph of a tool on a computer monitor. Participants were asked to “show how you would use the tool as if you were holding and using it” with the left hand. Four practice trials with feedback (using items different than in the task itself) were given at the start of the task. As per Rothi et al. (1991), if a participant gestured the action as if their hand was the tool (body-part-as-object error), they were reminded to “show how you would use the tool as if you were actually holding it in your hand”. The first of these errors was corrected and the participant was permitted a second try (for precedent, see Buxbaum et al., 2014; Garcea, Stoll, et al., 2019). Two items were determined to be outliers and were removed from the analysis of PTU in this NTC dataset (see Watson & Buxbaum, 2015). To maintain consistency with a prior group-level lesion-disconnectome mapping study Garcea et al., 2020, we retained these items in the PTU data of the LCVA group.

#### 2.3.2 Meaningless Gesture Imitation (MGI)

Participants were presented with videos of an experimenter performing 14 novel gestures and were instructed to imitate the gesture. Gestures were presented twice on each trial; during the first presentation, participants were instructed to watch the gesture in its entirety; at the beginning of the second presentation a sound was presented cueing participants to begin gesturing. The 14 novel gestures were developed to maintain similar spatio-motor characteristics of tool use gestures (e.g., plane of movement; joints moved; hand posture) but were designed such that the movement was meaningless (e.g., see Buxbaum et al., 2014). Fifty-one of the 58 LCVA participants took part in a version of the task with 10 meaningless actions, which were identical to 10 of the 14 meaningless actions used in the remaining 7 participants. To ensure there was no difference in MGI performance as a function of the number of trials in the task, we compared the mean hand posture accuracy of the 7 participants who imitated 14 gestures (mean = 0.6495; standard deviation = 0.1999) to the mean accuracy of the 51 participants who imitated 10 gestures (mean = 0.6486; standard deviation = 0.1970) and found no difference in performance (*t* < 1). All 17 NTC participants who performed the MGI task took part in the version of the task with 14 meaningless actions; the maximum number of trials for each NTC participant were used in the analysis of MGI performance.

#### 2.3.3 Coding of Action Data

Gestures were recorded with a digital camera and scored offline by two trained, reliable coders (Cohen’s Kappa score = 94%) who also demonstrated inter-rater reliability with previous coders in the Buxbaum lab (Cohen’s Kappa > 85%; see e.g., Buxbaum et al., 2005). Both tasks were coded using a portion of the detailed praxis scoring guidelines used in our previous work (see Buxbaum et al., 2003; Buxbaum et al., 2005; Watson & Buxbaum, 2015). In the PTU task, each gesture was given credit for semantic content unless a participant performed a recognizable gesture appropriate for another tool. Only gestures that were given credit for semantic content were scored on other action dimensions (e.g., spatio-temporal hand posture errors). Across both PTU and MGI tasks, hand posture errors were assigned if the shape or movement trajectory of the hand and/or wrist was flagrantly incorrect, or if the hand or finger was used as part of the tool (i.e., body-part-as-object error, Buxbaum et al., 2005; Watson & Buxbaum, 2015). This allowed us to investigate hand posture errors in the context of producing meaningful and meaningless actions, and to isolate unique variance associated with errors in either task. We used hand posture errors as the principal dependent variable because it can be measured across MGI and PTU tasks, and because prior work has demonstrated there is sensitivity in hand posture errors to measure abnormal action production in participants with limb apraxia (Garcea, Stoll, et al., 2019; Watson & Buxbaum, 2015). Our coding of hand posture errors in the MGI task did not differentiate between hand and finger imitation errors as in several prior investigations (e.g., see Achilles et al., 2019; Goldenberg & Karnath, 2006; Lesourd et al., 2018; Tessari et al., 2021).

### 2.4 Identification of Disproportionate Impairments Across PTU and MGI Tasks

For each participant, we computed the proportion of correct hand posture trials for the PTU and MGI tasks separately. Consistent with past findings in our lab (Buxbaum et al., 2014), there was a significant relation between performance in the two tasks (r(56) = 0.42, *p* < .01). We then identified participants with abnormal behavior across two independent analyses to ensure that effects were robust across cohorts and statistical methods.

#### 2.4.1 First-level Analysis: Linear Regression

The first-level analysis was restricted to participants with smaller lesions only. We rank ordered participants by total lesion volume and performed a median split, thereby identifying 29 participants with relatively smaller lesions. We then used regression residuals with this group of 29 to identify participants whose performance on PTU was disproportionately worse than expected based on their MGI performance, and vice versa. Residuals were then studentized to identify participants who performed disproportionately worse on PTU relative to MGI, or MGI relative to PTU. Studentized residuals were calculated by dividing each participant’s residual score by an estimate of the standard deviation excluding a given participant’s residual score. Given that we were interested in negative-going residual scores, we defined disproportionately worse performance using a one-tailed studentized *t*-score (*t*(28) = −1.70, *p* < .05).

### 2.4.2 Second-level Analysis: Single Test Dissociation

For each LCVA participant we performed two single Bayesian differences tests (Crawford et al., 2010)^1^. Each test quantified the degree to which the LCVA participant’s score on the PTU or MGI task dissociated from the control sample using Bayesian methods to estimate the probability that a member of the control group would obtain a score lower than the LCVA participant. In addition, a point estimate of the effect size of each LCVA participant’s performance relative to the control sample was calculated, along with 95% one-tailed confidence intervals and a one-tailed probability value.

We inspected the maximally disconnected subgraph of participants only if their performance differences were significant across both first- and second-level analyses.

### 2.5 Whole-brain Structural Disconnection Analysis

#### 2.5.1 Quantifying Whole-brain Structural Disconnection

For each participant we derived a whole-brain disconnectome, representing structural disconnection among cortical and subcortical nodes. Structural connectivity was inferred on the basis of stroke lesion location using a cohort of normal diffusion scans from The Human Connectome Project (HCP) database (for methodological detail, see Greene et al., 2019; for prior work in the investigation of apraxia severity, see Garcea et al., 2020). First, each LCVA participant’s stroke lesion was drawn on their corresponding native T1-weighted structural MRI volume and then normalized to a custom T1-weighted template constructed from 40 participants of the Human Connectome Project (HCP; Greene et al., 2018) using cost function masking in ANTS (Avants et al., 2008). We then parcellated each participant’s T1 structural volume using the Lausanne 2008 atlas (Daducci et al., 2012; Hagmann et al., 2008). The Lausanne atlas contains 130 nodes distributed throughout the right and left hemispheres; for the purpose of our analysis, we removed the cerebellum, resulting in 128 cortical and subcortical nodes (64 nodes in each hemisphere).

Structural connectivity between a given node pair was estimated for each of 210 neurotypical participants’ shortest path tractography derived from the diffusion scans of the HCP dataset. Then, each LCVA participant’s lesion was projected into these scans and the impact of the lesion on the shortest path tractography was estimated. Specifically, the percent loss in structural connectivity between any two nodes was calculated as the proportion of the shortest path tractography in the HCP dataset intersecting the lesion relative to the total shortest path probability shared between any given node pair in the HCP dataset (for methodological detail, see Greene et al., 2019). The result is a 128×128 disconnectome map, representing node-to-node (hereafter, edges) structural disconnection inferred on the basis of LCVA lesion location for each participant. Disconnection values were continuous, ranging from 1 (complete disconnection) to 0 (no disconnection). See Figure 1 for a visualization of each LCVA participant’s maximally disconnected subgraph.

#### 2.5.2 Calculating Maximally Disconnected Subgraphs

Because the full disconnectome can have many node pairs exhibiting negligible connectivity loss (rendering visualization and interpretation challenging), we reduce the dimensionality of each LCVA participant disconnectome by extracting a maximally disconnected subgraph. The subgraph is extracted by initializing a subgraph with the nodes that share the edges with the greatest structural connectivity loss. Additional nodes are added such that the cumulative amount or weight of the connectivity loss is maximized amongst the nodes; this procedure identifies a maximally disconnected subgraph when the change in the weight contributed by each additional node added reaches asymptote. The maximally disconnected subgraph contains regions and edges with the greatest shared structural connectivity loss induced by each LCVA lesion (see Greene et al., 2019). This algorithm was applied to the full disconnectome of the participants whose performance was identified as disproportionately worse for MGI or PTU.

### 3.1 Results

Plotted in Figure 2 is the mean performance in the PTU and MGI tasks for both LCVA and NTC groups.

**Figure 2.**
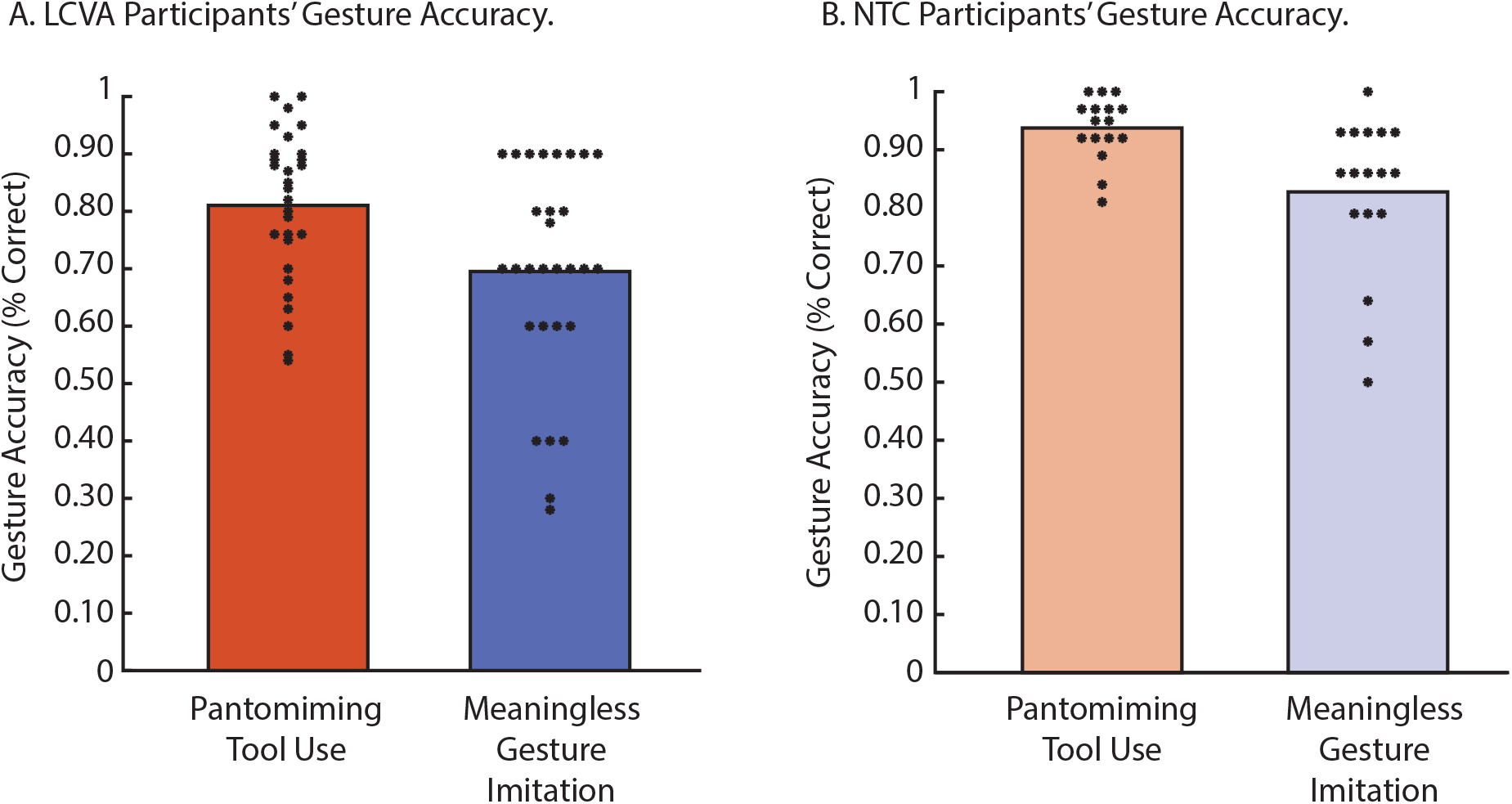
LCVA and NTC participants’ tool use pantomiming and meaningless gesture imitation accuracy scores. The mean accuracy PTU and MGI tasks are represented as bar plots separately for LCVA participants (A) and NTC participants (B). Each asterisk represents an individual participant’s accuracy value averaged across items.

### 3.2 Single-Case Analysis of PTU and MGI Performance

#### 3.2.1 Disproportionately Worse PTU Performance Compared to MGI: S20

S20 was identified as exhibiting relatively poor performance in the PTU task (S20’s mean = 0.55) when scores on this task were regressed against MGI performance (S20’s mean = 0.90) in the first-level analysis *t*(28) = −2.45, *p* < .001. The results of the second-level analysis indicated that 0.001% of control participants were estimated to score below S20’s PTU score. The difference between S20 and control participants was significant (*p* < .001) with an effect size of −6.98 (see Table 2A). In contrast, S20’s MGI performance was within control range: 57.16% of NTCs were estimated to score below S20’s MGI score (see Table 2B).

**Table 2.** Relative performance differences between PTU and MGI tasks in LCVA participants. **A.** Participant S20 was identified as performing disproportionately worse in the PTU task in relation to relatively normal performance in the MGI task. **B.** Participant S01 was identified as performing disproportionately worse in the MGI task in relation to relatively normal performance in the PTU task. *Abbreviations*. PTU HP %, Pantomiming tool use hand posture accuracy; MGI HP %, meaningless gesture imitation hand posture accuracy; CI, confidence interval.

**A. Analysis of PTU HP Performance, Controlling for MGI HP Performance.**
| LCVA Participant | Total Lesion Volume | Gesture Score (%) |  | First-Level Analysis | Second Level Analysis of PTU Performance |  |  |  |  |
| --- | --- | --- | --- | --- | --- | --- | --- | --- | --- |
|  |  | PTU HP | MGI HP | PTU ~ MGI | Estimated % of controls scoring lower than the LCVA participant |  | Estimated effect size (Z<sub>cc</sub>) |  | Single Bayes Significance Testing |
|  |  |  |  |  | Point Estimate | 95% Lower CI | Point Estimate | 95% Lower CI |  |
| S20 | 32003 | 0.55 | 0.90 | *t*(28) = -2.45, *p* < .001 | 0.001 | 0 | -6.98 | -9.04 | *p* < 0.001 |
| S01 | 337 | 0.95 | 0.40 | *t*(28) = 1.43, *p* = n.s. | 57.16 | 40.94 | 0.19 | -0.23 | *p* = n.s. |

**B. Analysis of MGI HP Performance, Controlling for PTU HP Performance.**
| LCVA Participant | Total Lesion Volume | Gesture Score (%) |  | First-Level Analysis | Second-Level Analysis of MGI Performance |  |  |  | Single Bayes Significance Testing |
| --- | --- | --- | --- | --- | --- | --- | --- | --- | --- |
|  |  | MGI HP | PTU HP | MGI ~ PTU | Estimated % of controls scoring lower than the LCVA participant |  | Estimated effect size (Z<sub>cc</sub>) |  |  |
|  |  |  |  |  | Point Estimate | 95% Lower CI | Point Estimate | 95% Lower CI |  |
| S01 | 337 | 0.40 | 0.95 | *t*(28) = -1.88, *p* < .05 | 0.41 | 0.002 | -3.11 | -4.06 | *p* = 0.004 |
| S20 | 32003 | 0.90 | 0.55 | *t*(28) = 1.60, *p* = n.s. | 70.35 | 54.87 | 0.56 | 0.12 | *p* = n.s. |

#### 3.2.2 Disproportionately Worse MGI Performance Compared to PTU

S01 was identified as exhibiting relatively poor performance in the MGI task (S01’s mean = 0.40) when scores on this task were regressed against PTU performance (S01’s mean = 0.95) in the first-level analysis *t*(28) = −1.88, *p* < .05. The results of the second-level analysis indicated that 0.41% of control participants were estimated to score below S01’s MGI score. The difference between S01 and NTCs was significant (*p* < .01) with an effect size of −3.11 (see Table 2B). In contrast, S01’s PTU performance was within control range: 70.35% of NTCs were estimated to score below S01’s PTU score (see Table 2A). For the full results of PTU and MGI first- and second-level analyses, see, respectively, Supplemental Tables 2A and 2B.

### 3.3 Single-Case Analysis of Maximally Disconnected Subgraph

#### 3.3.1 S20’s Lesion Location and Maximally Disconnected Subgraph

S20’s lesion site included portions of the left inferior parietal lobule (supramarginal gyrus and angular gyrus), superior and middle temporal gyri, middle and superior occipital cortex, and post-central gyrus (see Figure 3A and Table 3). S20’s maximally disconnected subgraph included disconnectivity among pre- and post-central gyri, inferior parietal lobule (supramarginal and angular gyri), superior parietal lobule, cuneus, precuneus, and superior, middle, and inferior temporal gyri. Disconnection extended inferiorly to include ventral temporal cortex (fusiform gyrus; lingual gyrus) and lateral occipital cortex, and anteriorly to include portions of inferior and middle frontal gyri (see Figure 3B; see Supplemental Table 3 for an ROI-to-ROI rendering of S20’s maximally disconnected subgraph). A node-level rendering of S20’s maximally disconnected subgraph is represented in Figure 3C.

**Figure 3.**
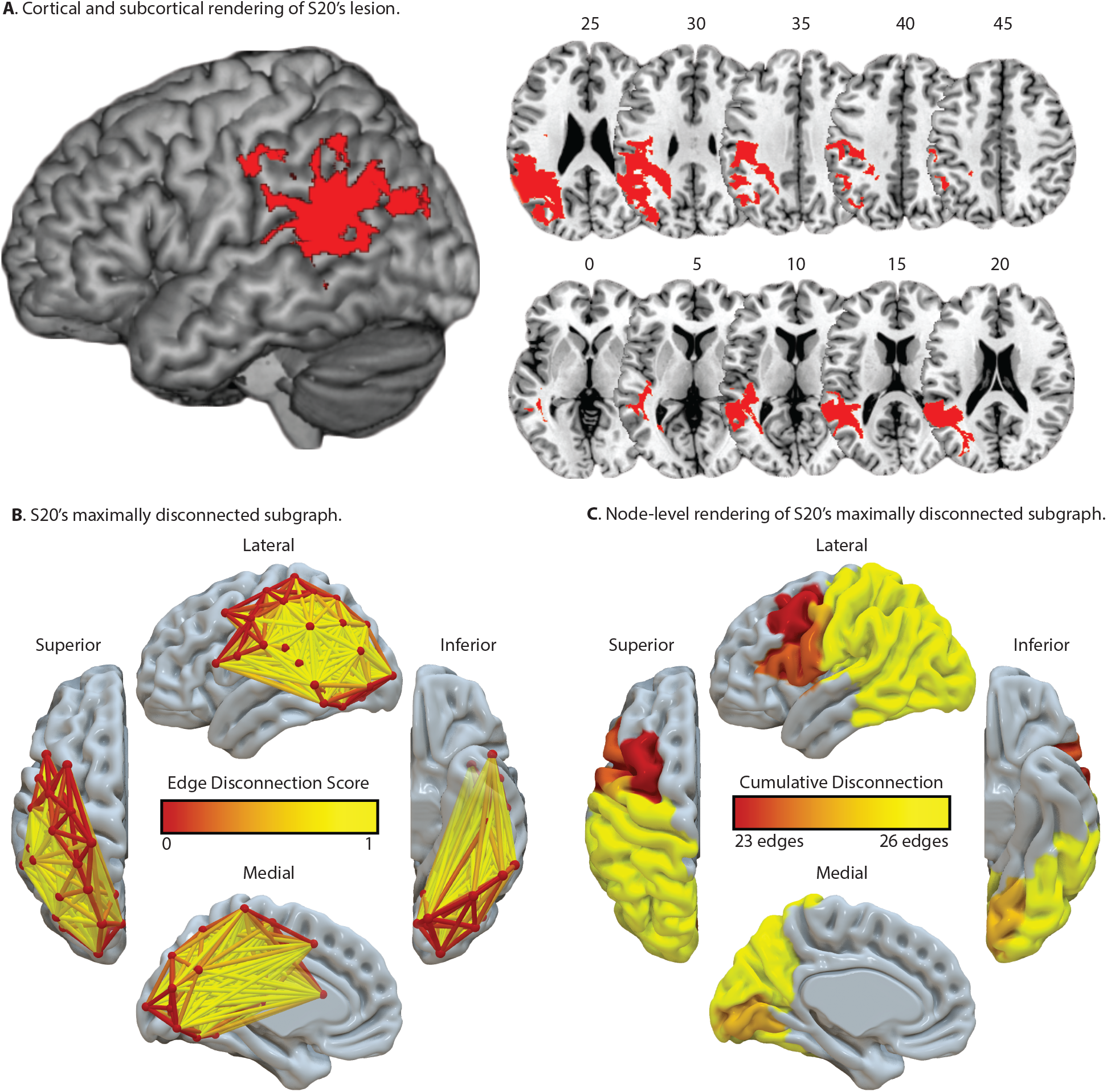
Visualization of S20’s lesion location and maximally disconnected subgraph. **A.** Cortical and subcortical rendering of S20’s lesion identifies the left inferior parietal lobule (supramarginal gyrus; angular gyrus), superior and middle temporal gyri, left postcentral gyrus, and middle and superior occipital cortices. Axial slices and corresponding MNI Z-coordinate values are presented in volumetric space. **B.** Analysis of S20’s maximally disconnected subgraph identifies disconnection among posterior regions in parietal, occipital, and temporal cortices, which extends anteriorly to include portions of the middle and inferior frontal gyri. **C.** A node-level rendering of S20’s maximally disconnected subgraph, including a count of disconnected edges per node.

**Table 3.** Proportion of voxels lesioned in S20 in neuroanatomic regions derived from the AAL Atlas. Regions with at least 1% damage to all voxels are included.

| AAL Atlas Label | AAL Number | Mean Center of Mass (XYZ) |  |  | % of Voxels Lesioned |
| --- | --- | --- | --- | --- | --- |
| Left Supramarginal Gyrus | 63 | -53 | -37 | 30 | 40.83 |
| Left Angular Gyrus | 65 | -46 | -58 | 31 | 35.99 |
| Left Superior Temporal Gyrus | 81 | -53 | -38 | 17 | 19.74 |
| Left Heschl's Gyrus | 79 | -39 | -23 | 10 | 13.19 |
| Left Middle Temporal Gyrus | 85 | -51 | -49 | 14 | 13.00 |
| Left Middle Occipital Cortex | 51 | -35 | -74 | 28 | 8.80 |
| Left Inferior Parietal Lobule | 61 | -47 | -38 | 41 | 8.01 |
| Left Superior Occipital Cortex | 49 | -22 | -69 | 26 | 3.41 |
| Left Postcentral Gyrus | 57 | -46 | -18 | 36 | 3.36 |

#### 3.3.2 S01’s Lesion Location and Maximally Disconnected Subgraph

S01’s lesion site included a portion of the left pre-central gyrus and adjacent white matter (see Figure 4A and Table 4). S01’s maximally disconnected subgraph included disconnectivity among pre- and post-central gyri, middle and superior frontal gyri, inferior and superior parietal lobule, and cuneus (see Figure 4B; see Supplemental Table 4 for an ROI-to-ROI rendering of S01’s maximally disconnected subgraph). A node-level rendering of S01’s maximally disconnected subgraph is represented in Figure 4C.

**Figure 4.**
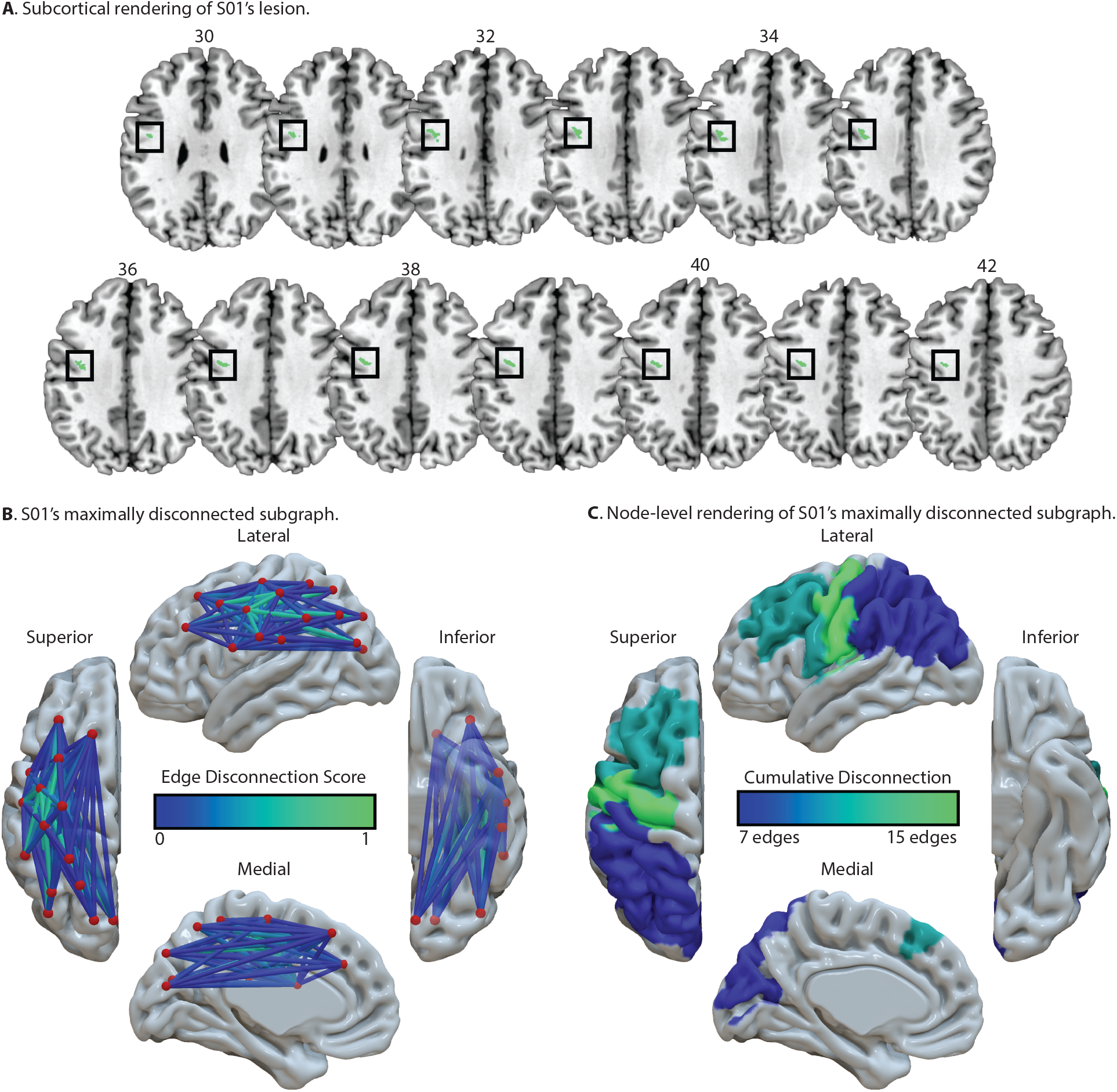
Visualization of S01’s lesion location and maximally disconnected subgraph. **A.** Cortical and subcortical rendering of S01’s lesion identifies minimal overlap with the left precentral gyrus. Axial slices and corresponding MNI Z-coordinate values are presented in volumetric space. **B.** Analysis of S01’s maximally disconnected subgraph identifies disconnection among inferior and superior parietal lobule, cuneus, pre- and post-central gyri, and superior and middle frontal gyri. **C.** A node-level rendering of S01’s maximally disconnected subgraph, including a count of disconnected edges per node.

**Table 4.** Proportion of lesioned voxels in S01. Anatomical labels are derived from the AAL Atlas. Only one region was identified in the left precentral gyrus.

| AAL Atlas Label | AAL Number | Mean Center of Mass (XYZ) |  |  | % of Voxels Lesioned |
| --- | --- | --- | --- | --- | --- |
| Left Precentral Gyrus | 1 | -40 | -4 | 35 | 0.31 |

### 4.1 General Discussion

The goal of this study was to investigate performance differences between PTU and MGI tasks at the single-case level to test the hypothesis that there are two subtypes of apraxia associated with distinct impairments in the conceptual versus production-related aspects of gesture, with distinct disconnectivity patterns among portions of the brain supporting praxis. Our first hypothesis was that worse performance in MGI relative to PTU would be associated with lesions to and disconnectivity among fronto-parietal nodes; our second hypothesis was that worse performance in PTU relative to MGI would be associated with lesions to and disconnectivity among temporo-parietal nodes. LCVA participants with small lesions were identified from a cohort of volunteers who performed the PTU and MGI tasks; we then identified participants who were disproportionately impaired at PTU given performance on MGI, and vice versa. Participant S20 was identified as exhibiting disproportionately worse performance with PTU compared to MGI. Analysis of S20’s lesion location revealed damage to and disconnection between the ventro-dorsal pathway and posterior temporal lobe, consistent with the hypothesis that structural connectivity between the left inferior parietal lobule and left posterior middle temporal gyrus is critical for tool use gesturing ability (Garcea et al., 2020). Moreover, S20’s disconnection extended into visual cortex to include the fusiform gyrus, lingual gyrus, and lateral occipital cortex, consistent with prior investigations demonstrating these regions are involved in tool recognition and use (Almeida et al., 2013; Garcea, Almeida, et al., 2019; Mahon et al., 2013; Watson et al., 2019). S01, in contrast, exhibited disproportionately worse performance with MGI compared to PTU. Analysis of S01’s lesion location and disconnection revealed damage to parietal, premotor, and frontal structures, revealing a pattern consistent with the dorso-dorsal pathway identified in prior group-level VLSM studies (Buxbaum et al., 2014; Goldenberg et al., 2007; Martin et al., 2016; Mengotti et al., 2013; Tessari et al., 2007; Weiss et al., 2016, for review, see Rumiati et al., 2009). Below, we discuss our findings and their implications for a neurocognitive model of praxis in the human brain.

### 4.2 Implications for Neurocognitive Models of Praxis

Cognitive models of praxis posit a distinction between a direct and an indirect route supporting skilled action (Buxbaum, 2001; Buxbaum & Randerath, 2018; Cubelli et al., 2000; Rothi et al., 1997; Rumiati & Humphreys, 1998). The direct route is argued to support the production of gestures by transforming current visual input into motor output, driven ‘bottom-up’ by the stimulus. In contrast, the indirect route interfaces current visual input with stored semantic knowledge for gesture production. The indirect route facilitates processing when gesturing a meaningful action (e.g., eating soup with a spoon) because the implementation of a meaningful gesture is supported by semantic memory processes, including the retrieval of function knowledge (e.g., that a spoon is used for eating), the retrieval of hand posture knowledge (e.g., that a spoon is grasped and manipulated with a specific configuration of the hand and fingers), and the retrieval of the visual appearance of a gesture (e.g., that a spoon moves along a trajectory from a bowl to the mouth when eating soup). Whereas performance in meaningless gesture imitation is sensitive the contributions of the direct route, as stored action knowledge, by hypothesis, is not required to imitate a meaningless gesture, pantomime of tool use is hypothesized to reflect the integrity of the indirect route (e.g., see Buxbaum, 2017; Rothi et al., 1991; Rumiati et al., 2010). Differential performance across meaningless gesture imitation and pantomime of tool use tasks is consistent with the notion of a direct and indirect route, respectively, supporting skilled action (e.g., see Rumiati et al., 2010; Tessari et al., 2007).

Given the partially overlapping neural architecture of the two routes for action posited by the 2AS+ model, it is unsurprising that there were correlations in PTU and MGI performance, as LCVA participants with moderate to large lesions often demonstrate deficits associated with both pathways. However, our approach identified participants who presented with a distinct cognitive profile and a relatively focal lesion and structural disconnection pattern. For instance, S20 had a temporo-parietal lesion and was disproportionately worse at PTU (consistent with a ‘representational’ or ‘conceptual’ apraxia subtype; for discussion, see Buxbaum & Kalenine, 2021), whereas S01 had a left precentral gyrus lesion resulting in disconnection in superior parietal areas, and was disproportionately worse at MGI (consistent with a ‘production’ apraxia subtype; for discussion, see Buxbaum & Kalenine, 2021). The behavioral patterns resulting from these lesion loci are consistent with the claim that the ventro-dorsal and dorso-dorsal pathways support distinct aspects of skilled action production. Moreover, the results of S01’s structural disconnection analysis are especially remarkable given their lesion size and performance differences. S01’s structural disconnectivity results are consistent with prior findings that praxis ability is supported by longitudinal tracts connecting the left inferior parietal lobe with frontal sites, including the superior longitudinal fasciculus (SLF; see Ramayya et al., 2010). For example, Vry and colleagues (2015) demonstrated that SLF fiber bundles connecting parietal and frontal regions—including dorsal premotor cortex—supports the imitation of meaningless hand gestures. Interestingly, prior work indicates that the SLF is critical for pantomime of tool use, as lesions overlapping with the SLF are associated with reduced tool use pantomiming ability (e.g., see Garcea, Stoll, et al., 2019; Watson & Buxbaum, 2015). Thus, it will be critical for future work to determine if deficits in meaningless gesture imitation and pantomime of tool use result from damage to dissociable fiber bundles of the SLF.

A growing body of research has demonstrated that there is increased task-based functional connectivity between the left supramarginal gyrus and the left posterior middle temporal gyrus when neurotypical participants pantomime the use of tools (Garcea et al., 2018; Hutchison & Gallivan, 2018; Vingerhoets & Clauwaert, 2015); this finding has also been observed in resting state functional connectivity studies (Hutchison et al., 2014; Simmons & Martin, 2012). Moreover, it has recently been shown that the degree of resting state functional connectivity reduction between left inferior parietal lobule and left posterior middle temporal gyrus was associated with the severity of reduced tool use pantomiming performance (Watson et al., 2019). Our theoretic framework hypothesizes that successful retrieval of hand posture knowledge for tool use pantomiming is supported in part by functional interactions between the posterior middle temporal gyrus and the left inferior parietal lobule (e.g., see Garcea & Buxbaum, 2019; Garcea et al., 2020; Watson et al., 2019). By hypothesis, the left posterior middle temporal gyrus is involved in the retrieval of the visual appearance of a tool use action, and the left inferior parietal lobule aggregates visual attributes of a tool and its associated action (e.g., that a hammer is swung along a particular trajectory to pound in a nail) with fronto-parietal visuomotor circuits that support the programming and implementation of tool use actions (for discussion, see Finkel et al., 2018; Frey, 2007; Kroliczak et al., 2016). Successful tool use pantomiming is the result of functional interactions among temporal, parietal, and frontal regions working in concert to interface long-term semantic memory of actions and tools with online processes that enable skilled motor production. The result from our analysis of S20’s maximally disconnected subgraph provides evidence at the single-case level that disconnection between inferior parietal and lateral posterior temporal regions is associated with reduced tool use pantomiming ability, suggesting that S20’s tool use pantomiming deficit may derive from an impairment in the retrieval of hand posture knowledge when pantomime of tool use actions.

### 4.3 Limitations and Considerations

It’s important to note that we did not compute a double dissociation metric commonly used in case study investigations (Crawford & Garthwaite, 2006; Crawford et al., 2010). Thus, we cannot conclude that our single-case results constitute a double dissociation between cognitive mechanisms or between neuroanatomic substrates. Nevertheless, consider the performance of S01, whose MGI accuracy (0.40) was disproportionately lower in relation to their PTU accuracy (0.95) despite having the smallest lesion of the cohort (337 voxels). S01’s structural disconnection was restricted to fronto-parietal regions, avoiding posterior temporal lobe structures hypothesized to support pantomime of tool use. Though speculative, this individual may represent a ‘production’ subtype of apraxia, with a lesion site and structural disconnectivity pattern that localizes to the dorso-dorsal pathway (for discussion, see Buxbaum & Kalenine, 2021).

A second consideration is that structural disconnection was inferred using shortest path tractography. Though this method has demonstrated sensitivity to corticospinal tract damage in relation to weakened contralateral hand grip strength (Greene et al., 2019), this metric provides a marker of cortical-to-cortical disconnection, and is therefore agnostic to the specific white matter tracts that are damaged. Thus, it will be important for future studies to identify the appropriate structural connectivity method (i.e., inferred or computed from diffusion imaging data) in addition to the fiber tracking algorithms used to quantify structural disconnection (e.g., probabilistic, deterministic, shortest path tractography) when investigating structure-function relations in participants with brain damage due to stroke or other neurological diseases (e.g., see Bi et al., 2015; Forkel et al., 2021; Gleichgerrcht et al., 2017; Griffis et al., 2019; Kuceyeski et al., 2016; Rosenzopf et al., 2022; Salvalaggio et al., 2020; Yeh et al., 2013).

Finally, although prior work has inspected inter-hemispheric functional connectivity associated with recovery of language and motor function (e.g., see Carter et al., 2010; Siegel et al., 2016; for review, see Grefkes & Fink, 2014), the cognitive and neural mechanisms associated with recovery of praxis function remain poorly understood. Moving forward, it will be critical to investigate the relation among inter-hemispheric structural connectivity, functional connectivity, and apraxia recovery, as recent work demonstrated that greater inter-hemispheric resting state functional connectivity was associated with less severe tool use pantomiming impairment in persons with chronic apraxia (Watson et al., 2019).

### 4.4 Conclusions

The neuroanatomy supporting meaningless gesture imitation and pantomime of tool use is distributed in functionally and computationally distinct pathways within the human dorsal visual pathway. Our approach combines cognitive performance and lesion location to identify deviations across gesturing tasks at the single-case level, which is used to test hypotheses of lesion loci and structural disconnectivity among regions critical for skilled action production. Our results suggest that impairments in conceptual or production aspects of skilled action are associated with distinct patterns of lesion-induced structural disconnection, consistent with the ventro-dorsal pathway supporting key aspects of tool use pantomiming, and the dorso-dorsal pathway supporting key aspects of meaningless gesture imitation. Furthermore, this method may be useful for the identification of subtypes of apraxia associated with distinct behavioral deficits, lesion loci, and structural disconnectivity patterns. The approach we have used can be tailored to investigate the cognitive and neuroanatomic substrates of other neuropsychological phenomena, including aphasia, optic ataxia, alexia, and visuospatial neglect.

## Acknowledgements

We thank Cortney Howard and Veronica Kreter for coding participants’ gestures, and Austin Wild, Olu Faseyitan, Branch Coslett, and Clint Greene for help with lesion segmentation. Preparation of this manuscript was supported by a Moss Rehabilitation Research Institute/University of Pennsylvania postdoctoral training fellowship (NIH 5T32HD071844-05), and by NIH grant R01 NS099061 to L.J.B.

## Author Information

The authors declare no competing financial interests. Correspondence should be addressed to F.E.G.

## Data Availability Statement

In compliance with the guidelines of the Institutional Review Board of Albert Einstein Healthcare Network, all participants gave informed consent and were compensated for travel expenses and participation. The informed consents obtained did not include permission to make data publicly available. Accordingly, the conditions of our ethical approval do not permit anonymized data to be publicly archived. To obtain access to the data, individuals should contact the corresponding author. Requests for data are assessed and approved by the Institutional Review Board of Einstein Healthcare Network.

## Supplemental Online Materials

**Supplemental Figure 1.**
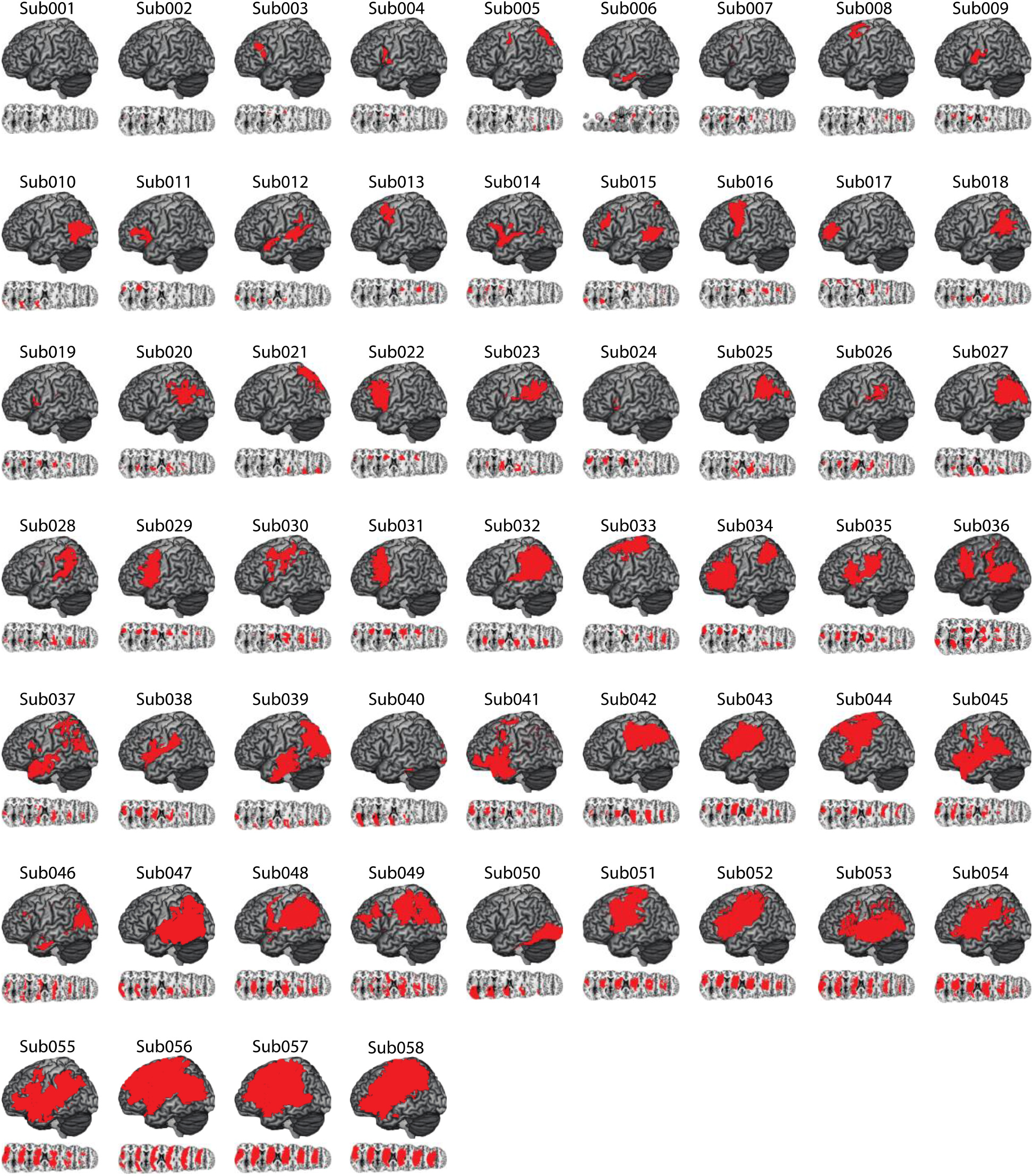
Distribution of Cortical and Subcortical Damage in Each LCVA Participant. A depiction of the 58 LCVA participants’ lesions presented on the ch2bet template. Lesions are projected on the cortical surface but include subcortical voxels using a 12-voxel search depth. Axial slices begin at the MNI coordinate [0,0,0] and increase in 10-mm increments superiorly.

**Supplemental Table 1.** Demographic information and lesion volume for LCVA participants with relatively larger lesions. *Abbreviations*. PTU, Pantomime of tool use to the sight of objects; MGI, meaningless gesture imitation; HP, hand posture accuracy.

| LCVA Participant | PTU HP (%) | MGI HP (%) | Months Post Stroke Onset | Lesion Volume (1mm<sup>3</sup>) | Years of Education | Gender | Age |
| --- | --- | --- | --- | --- | --- | --- | --- |
| S30 | 0.74 | 0.60 | 9 | 56281 | 9 | F | 66 |
| S31 | 0.50 | 0.40 | 7 | 61198 | 12 | F | 71 |
| S32 | 0.56 | 0.40 | 9 | 62204 | 18 | F | 32 |
| S33 | 0.85 | 0.78 | 184 | 62530 | 13 | F | 60 |
| S34 | 0.98 | 1.00 | 6 | 64375 | 12 | M | 45 |
| S35 | 0.79 | 0.70 | 8 | 71022 | 16 | F | 60 |
| S36 | 0.26 | 0.50 | 7 | 71905 | 18 | M | 67 |
| S37 | 0.73 | 0.70 | 14 | 76301 | 18 | M | 64 |
| S38 | 0.83 | 0.50 | 84 | 80020 | 13 | F | 47 |
| S39 | 0.89 | 0.71 | 151 | 80532 | 16 | F | 55 |
| S40 | 0.80 | 0.60 | 47 | 82964 | 14 | M | 53 |
| S41 | 0.87 | 0.64 | 30 | 87120 | 18 | M | 76 |
| S42 | 0.85 | 0.70 | 10 | 88046 | 13 | M | 31 |
| S43 | 0.83 | 0.78 | 6 | 92744 | 12 | M | 48 |
| S44 | 0.90 | 0.86 | 17 | 93628 | 16 | F | 37 |
| S45 | 0.93 | 0.50 | 68 | 94536 | 12 | F | 45 |
| S46 | 0.80 | 0.50 | 161 | 96196 | 13 | M | 60 |
| S47 | 0.87 | 0.40 | 17 | 102522 | 16 | F | 38 |
| S48 | 0.60 | 0.70 | 174 | 115118 | 16 | F | 54 |
| S49 | 0.63 | 0.50 | 19 | 117809 | 12 | F | 41 |
| S50 | 0.83 | 0.70 | 9 | 128897 | 12 | F | 54 |
| S51 | 0.66 | 0.70 | 32 | 136576 | 12 | F | 46 |
| S52 | 0.83 | 0.90 | 112 | 166393 | 13 | M | 57 |
| S53 | 0.55 | 0.00 | 22 | 171128 | 18 | M | 51 |
| S54 | 0.62 | 0.60 | 31 | 179606 | 15 | M | 57 |
| S55 | 0.70 | 0.70 | 31 | 200079 | 12 | M | 61 |
| S56 | 0.39 | 0.40 | 12 | 225021 | 16 | M | 64 |
| S57 | 0.75 | 0.50 | 172 | 258736 | 18 | M | 68 |
| S58 | 0.24 | 0.50 | 11 | 303310 | 12 | F | 65 |

**Supplemental Table 2.** Participant gesture scores and results from first- and second-level analyses. In the first-level analysis a median split was used to remove participants with relatively larger lesions, and raw gesture scores were entered in a regression analysis, resulting in a studentized *t*-score indicative of the extent to which performance on one task was disproportionately lower than performance on the other task. In the second-level analysis each participant’s score was compared to the mean and standard deviation of the control samples. Bayesian methods were used to estimate the probability th at the participant’s score would be observed in the control sample as well as the effect size (in units of standard deviation relative to the control sample). Bolded LCVA participants presented with disproportionately worse performance for PTU or MGI tasks across first-and second-level analyses; italicized LCVA participants presented with impaired performance for PTU and MGI tasks across first- and second-level analyses. *Abbreviations*: PTU ∼ MGI, pantomiming tool use hand posture accuracy controlling for meaningless gesture imitation hand posture accuracy; MGI ∼ PTU, meaningless gesture imitation hand posture accuracy controlling for pantomiming tool use hand posture accuracy; CI, confidence interval.

| A. Analysis of PTU HP Performance, Controlling for MGI HP Performance. |  |  |  |  |  |  |  |  |  |
| --- | --- | --- | --- | --- | --- | --- | --- | --- | --- |
| LCVA Participant | Total Lesion Volume | Gesture Score (%) |  | First-Level Analysis |  | Second Level Analysis of PTU Performance |  |  |  |
|  |  | PTU HP | MGI HP | PTU ~ MGI studentized *t*-values | Estimated % of controls scoring lower than the LCVA participant |  | Estimated effect size (*Z*<sub>CC</sub>) |  | Single Bayes Significance Testing |
|  |  |  |  |  | Point Estimate | 95% Lower CI | Point Estimate | 95% Lower CI |  |
| S01 | 337 | 0.95 | 0.40 | 1.43 | 57.16 | 40.94 | 0.19 | -0.23 | *p* = 0.43 |
| S02 | 1869 | 0.82 | 0.90 | -0.11 | 2.79 | 0.20 | -2.14 | -2.87 | *p* = 0.03 |
| S03 | 4095 | 0.95 | 0.60 | 1.18 | 58.05 | 41.82 | 0.21 | -0.21 | *p* = 0.42 |
| S04 | 5953 | 0.79 | 0.40 | 0.13 | 1.11 | 0.02 | -2.63 | -3.49 | *p* = 0.01 |
| S05 | 9489 | 0.80 | 0.70 | -0.08 | 1.46 | 0.05 | -2.47 | -3.31 | *p* = 0.01 |
| S06 | 14660 | 0.90 | 0.70 | 0.69 | 27.78 | 12.82 | -0.69 | -1.14 | *p* = 0.25 |
| S07 | 16186 | 0.65 | 0.60 | -1.15 | 0.007 | 0 | -5.15 | -6.69 | *p* < 0.001 |
| S08 | 16977 | 0.89 | 0.90 | 0.44 | 20.86 | 9.15 | -0.86 | -1.33 | *p* = 0.41 |
| S09 | 16978 | 1.00 | 0.90 | 1.34 | 85.13 | 71.56 | 1.11 | 0.57 | *p* = 0.15 |
| S10 | 17706 | 0.98 | 0.80 | 1.20 | 73.50 | 57.80 | 0.66 | 0.20 | *p* = 0.26 |
| S11 | 18528 | 0.54 | 0.28 | -2.03 | 0.002 | 0 | -7.22 | -9.35 | *p* < 0.001 |
| S12 | 20105 | 1.00 | 0.60 | 1.61 | 85.13 | 71.56 | 1.11 | 0.57 | *p* = 0.15 |
| S13 | 20190 | 0.63 | 0.60 | -1.32 | 0.004 | 0 | -5.52 | -7.16 | *p* < 0.001 |
| S14 | 20790 | 0.76 | 0.80 | -0.45 | 0.41 | 0.002 | -3.14 | -4.13 | *p* = 0.004 |
| S15 | 23141 | 0.68 | 0.70 | -1.05 | 0.02 | 0 | -4.74 | -6.17 | *p* < 0.001 |
| S16 | 27004 | 0.87 | 0.70 | 0.45 | 12.55 | 3.87 | -1.23 | -1.77 | *p* = 0.13 |
| S17 | 27840 | 0.90 | 0.70 | 0.67 | 24.37 | 11.74 | -0.73 | -1.19 | *p* = 0.24 |
| S18 | 29052 | 0.76 | 0.90 | -0.55 | 0.40 | 0.001 | -3.15 | -4.14 | *p* = 0.004 |
| S19 | 30414 | 0.89 | 0.80 | 0.56 | 23.01 | 10.72 | -0.78 | -1.24 | *p* = 0.43 |
| **S20** | 32003 | 0.55 | 0.90 | **-2.45** | 0.001 | 0 | -6.98 | -9.04 | ***p* < 0.001** |
| S21 | 32684 | 0.70 | 0.30 | -0.54 | 0.04 | 0 | -4.29 | -5.59 | $p < 0.001$ |
| S22 | 33183 | 0.60 | 0.70 | -1.69 | 0.001 | 0 | -6.08 | -7.89 | $p < 0.001$ |
| S23 | 37091 | 0.93 | 0.70 | 0.88 | 41.07 | 25.79 | -0.24 | -0.65 | $p = 0.41$ |
| S24 | 44493 | 0.85 | 0.40 | 0.60 | 7.23 | 1.41 | -1.59 | -2.19 | $p = 0.07$ |
| S25 | 47442 | 0.88 | 0.70 | 0.49 | 14.38 | 4.91 | -1.14 | -1.65 | $p = 0.14$ |
| S26 | 48459 | 0.88 | 0.78 | 0.42 | 14.38 | 4.91 | -1.14 | -1.65 | $p = 0.14$ |
| S27 | 51399 | 0.76 | 0.90 | -0.55 | 0.41 | 0.002 | -3.14 | -4.13 | $p = 0.004$ |
| S28 | 51780 | 0.84 | 0.90 | 0.06 | 5.72 | 0.90 | -1.73 | -2.37 | $p = 0.06$ |
| S29 | 52416 | 0.75 | 0.90 | -0.66 | 0.24 | 0 | -3.39 | -4.44 | $p = 0.002$ |

**B.** Analysis of MGI HP Performance, Controlling for PTU HP Performance.
| LCVA Participant | Total Lesion Volume | Gesture Score (%) |  | First-Level Analysis |  | Second Level Analysis of MGI Performance |  |  |  |
| --- | --- | --- | --- | --- | --- | --- | --- | --- | --- |
| | | MGI HP | PTU HP | MGI ~ PTU studentized $t$-values | Estimated % of controls scoring lower than the LCVA participant | | Estimated effect size ($Z_{cc}$) | | Single Bayes Significance Testing |
|  |  |  |  |  | Point Estimate | 95% Lower CI | Point Estimate | 95% Lower CI |  |
| **S01** | 337 | 0.40 | 0.95 | **-1.88** | 0.41 | 0.002 | -3.11 | -4.06 | $p = 0.004$ |
| S02 | 1869 | 0.90 | 0.82 | 1.09 | 70.35 | 54.87 | 0.56 | 0.12 | $p = 0.29$ |
| S03 | 4095 | 0.60 | 0.95 | -0.70 | 6.53 | 1.25 | -1.64 | -2.24 | $p = 0.07$ |
| S04 | 5953 | 0.40 | 0.79 | -1.60 | 0.41 | 0.002 | -3.11 | -4.06 | $p = 0.004$ |
| S05 | 9489 | 0.70 | 0.80 | 0.04 | 19.58 | 8.52 | -0.91 | -1.37 | $p = 0.20$ |
| S06 | 14660 | 0.70 | 0.90 | -0.09 | 19.58 | 8.52 | -0.91 | -1.37 | $p = 0.20$ |
| S07 | 16186 | 0.60 | 0.65 | -0.31 | 6.53 | 1.25 | -1.64 | -2.24 | $p = 0.07$ |
| S08 | 16977 | 0.90 | 0.89 | 1.00 | 70.35 | 54.87 | 0.56 | 0.12 | $p = 0.30$ |
| S09 | 16978 | 0.90 | 1.00 | 0.88 | 70.35 | 54.87 | 0.56 | 0.12 | $p = 0.30$ |
| S10 | 17706 | 0.80 | 0.98 | 0.35 | 43.45 | 28.37 | -0.17 | -0.57 | $p = 0.43$ |
| S11 | 18528 | 0.28 | 0.54 | -2.18 | 0.07 | 0 | -3.99 | -5.17 | $p < 0.001$ |
| S12 | 20105 | 0.60 | 1.00 | -0.78 | 6.53 | 1.25 | -1.64 | -2.24 | $p = 0.07$ |
| S13 | 20190 | 0.60 | 0.63 | -0.28 | 6.53 | 1.25 | -1.64 | -2.24 | $p = 0.07$ |
| S14 | 20790 | 0.80 | 0.76 | 0.62 | 43.45 | 28.37 | -0.17 | -0.57 | $p = 0.43$ |
| S15 | 23141 | 0.70 | 0.68 | 0.20 | 19.58 | 8.52 | -0.91 | -1.37 | $p = 0.20$ |
| S16 | 27004 | 0.70 | 0.87 | -0.05 | 19.58 | 8.52 | -0.91 | -1.37 | $p = 0.20$ |
| S17 | 27840 | 0.70 | 0.90 | -0.09 | 19.58 | 8.52 | -0.91 | -1.37 | $p = 0.20$ |
| S18 | 29052 | 0.90 | 0.76 | 1.17 | 70.35 | 54.87 | 0.56 | 0.12 | $p = 0.30$ |
| S19 | 30414 | 0.80 | 0.89 | 0.45 | 43.45 | 28.37 | -0.17 | -0.57 | $p = 0.43$ |
| S20 | 32003 | 0.90 | 0.55 | 1.60 | 70.35 | 54.87 | 0.56 | 0.12 | $p = 0.30$ |
| S21 | 32684 | 0.30 | 0.70 | -2.13 | 0.09 | 0 | -3.84 | -4.99 | $p < 0.001$ |
| S22 | 33183 | 0.70 | 0.60 | 0.31 | 19.58 | 8.52 | -0.91 | -1.37 | $p = 0.20$ |
| S23 | 37091 | 0.70 | 0.93 | -0.12 | 19.58 | 8.52 | -0.91 | -1.37 | $p = 0.20$ |
| S24 | 44493 | 0.40 | 0.85 | -1.69 | 0.41 | 0.002 | -3.11 | -4.06 | $p = 0.004$ |
| S25 | 47442 | 0.70 | 0.88 | -0.06 | 19.58 | 8.52 | -0.91 | -1.37 | $p = 0.20$ |
| S26 | 48459 | 0.78 | 0.88 | 0.37 | 38.02 | 23.47 | -0.32 | -0.72 | $p = 0.38$ |
| S27 | 51399 | 0.90 | 0.76 | 1.17 | 70.35 | 54.87 | 0.56 | 0.12 | $p = 0.30$ |
| S28 | 51780 | 0.90 | 0.84 | 1.06 | 70.35 | 54.87 | 0.56 | 0.12 | $p = 0.30$ |
| S29 | 52416 | 0.90 | 0.75 | 1.19 | 70.35 | 54.87 | 0.56 | 0.12 | $p = 0.30$ |

**Supplemental Table 3.**
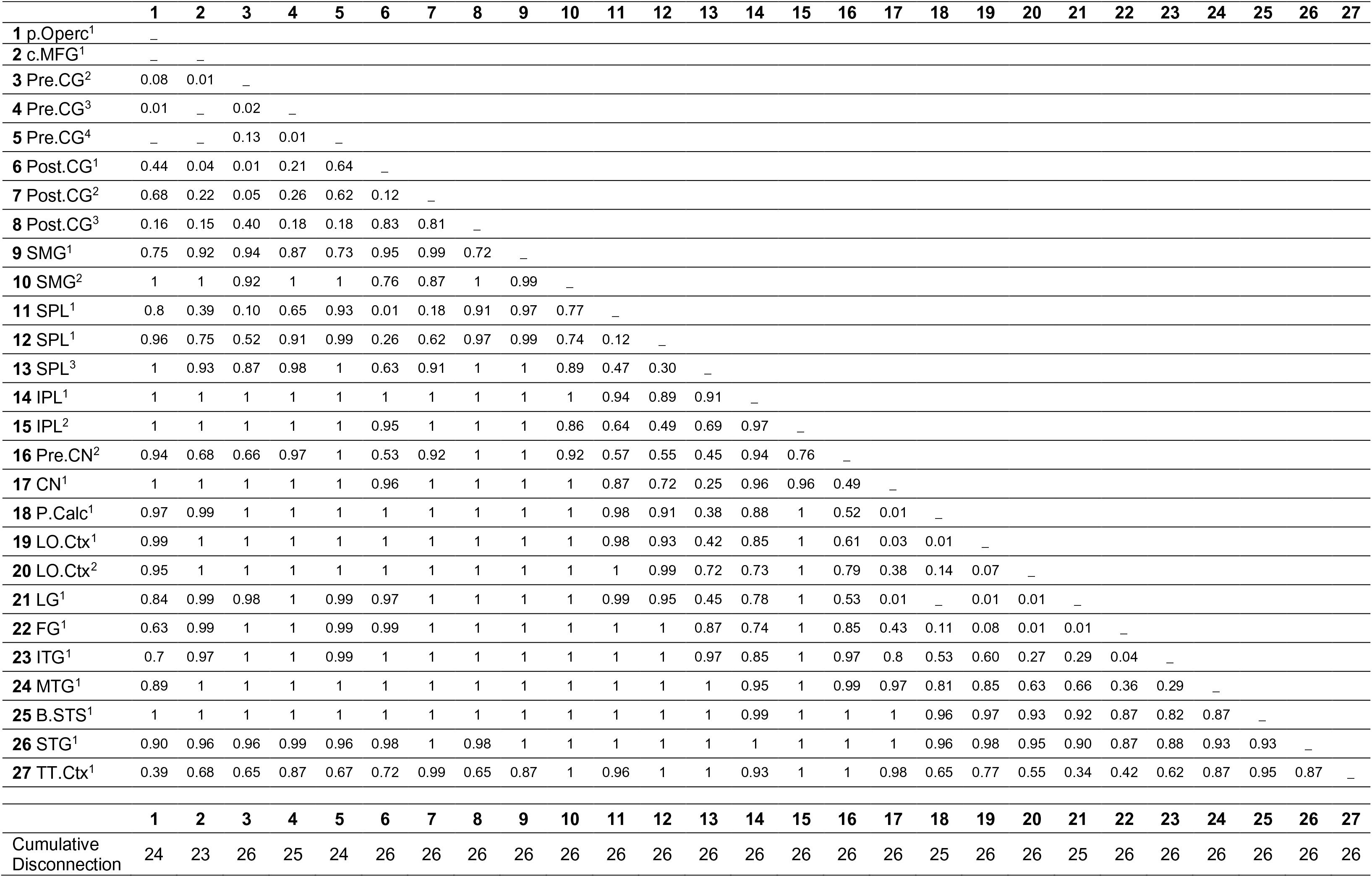
Maximally disconnected subgraph of S20. *Abbreviations*: (1) p.Operc, Pars Opercularis; (2) c.MFG, Caudal Middle Frontal Gyrus; (3:5) Pre.CG, Pre-central Gyrus; (6:8) Post.CG, Post-central Gyrus; (9:10) SMG, Supramarginal Gyrus; (11:13) SPL, Superior Parietal Lobule; (14:15) IPL, Inferior Parietal Lobule (Angular Gyrus); (16) Pre.CN, Precuneus; (17) CN, Cuneus; (18) P.Calc, Pericalcarine Cortex; (19:20) LO.CTX, Lateral Occipital Cortex; (21) LG, Lingual Gyrus; (22) FG, Fusiform Gyrus; (23) ITG, Inferior Temporal Gyrus; (24) MTG, Middle Temporal Gyrus; (25) STG, Superior Temporal Gyrus; (26) B.STS, Base of the Superior Temporal Sulcus; (27) TT.Ctx, Transverse Temporal Cortex. Numbers in superscript denote anatomical subregion from the Lausanne Atlas.

**Supplemental Table 4.**
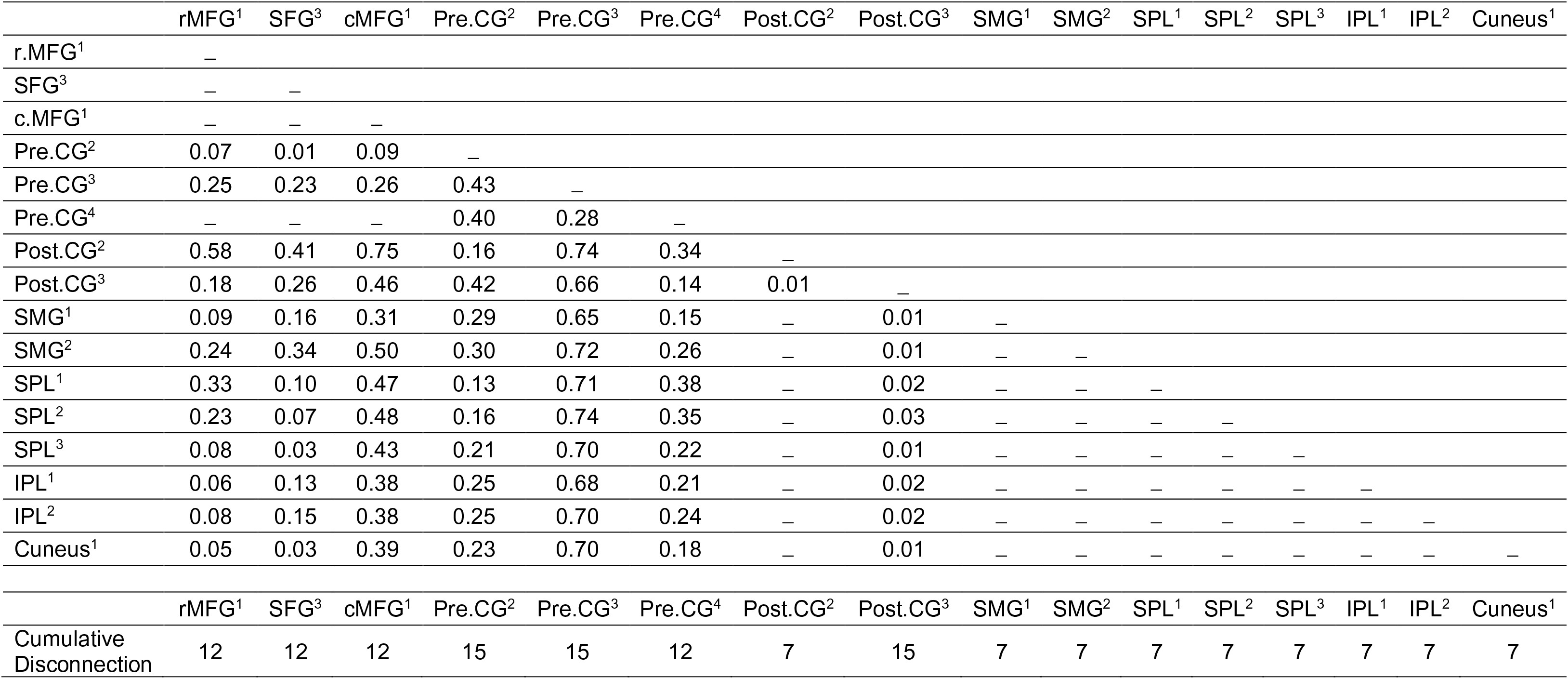
Maximally disconnected subgraph of S01. *Abbreviations*: r.MFG^1^, Rostral Middle Frontal Gyrus; SFG^3^, Superior Frontal Gyrus; c.MFG^1^, Caudal Middle Frontal Gyrus; Pre.CG, Pre-central Gyrus; Post.CG, Post-central Gyrus; SMG, Supramarginal Gyrus; SPL, Superior Parietal Lobule; IPL, Inferior Parietal Lobule (Angular Gyrus). Numbers in superscript denote anatomical subregion from the Lausanne Atlas.

## Footnotes

1 The SingleBayes_ES program can be downloaded at https://homepages.abdn.ac.uk/j.crawford/pages/dept/Single_Case_Effect_Sizes.htm

